# Uncovering Invariant Representations in Functional Neuroimaging with Deep Metric Learning

**DOI:** 10.1101/2023.09.17.558181

**Authors:** Arunesh Mittal, Xiaoxiao Sun, John Paisley, Paul Sajda

## Abstract

With the increasing ability to record neuroimaging with higher spatial and temporal resolution, there is a growing need for methods that reduce these high-dimensional representations into latent low-dimensional structures that are discriminative and/or predictive of behavior, disease, or in general experimental context. We propose a metric learning framework to extract meaningful latent structures from high-dimensional fMRI data. This method learns the latent embeddings that reduce the intra-group variability while maximizing the inter-group variability. In addition, our method leverages advances in few-shot learning approaches to adapt to small sample-size fMRI datasets, allowing one to learn the latent structure from just a few samples per context. We evaluate our work on two publicly available fMRI datasets and report superior results compared to popular alternative approaches such as Principal Component Analysis (84.7% vs. 60%; 21.8% vs. 8.3%). We provide the Python code as open-source at Github.

## 1. Introduction

Central to the field of neuroimaging is the goal of meaningfully connecting neural activity with complex cognitive behaviors and clinical conditions (Fox & Greicius, 2010; Reinhardt et al., 2010; Urai et al., 2022). Challenging is that neural activity is only partially observable and inferences about how this activity links to behavior are wrought with confounds and limited statistical power (Varoquaux & Thirion, 2014; Cremers et al., 2017; Marek et al., 2022). This has led to new efforts for developing correlative and predictive machine learning methods that yield interpretable (i.e., non-black-box) results given limited data set size and high dimensional data. This is particularly true in the analysis of functional magnetic resonance imaging (fMRI), where multivariate machine learning methods have begun to replace classic univariate methods for analysis (Lizier et al., 2011; Abraham et al., 2014; Khosla et al., 2019; Nielsen et al., 2020; Kohoutova et al., 2020). Though such machine learning methods have yielded promising results in terms of correlating and/or predicting behavior or clinical condition, interpretability is still challenging since it requires understanding the uncovered latent low-dimensional structure that drives the decoding model (Chen et al., 2013; Vu et al., 2020). In addition, the presence of both intra-group variability and inter-group variability makes assigning variance in the latent space challenging. For example, a goal is typically to optimize inter-class variability while reducing intra-class variability (See Figure 1).

**Figure 1.**
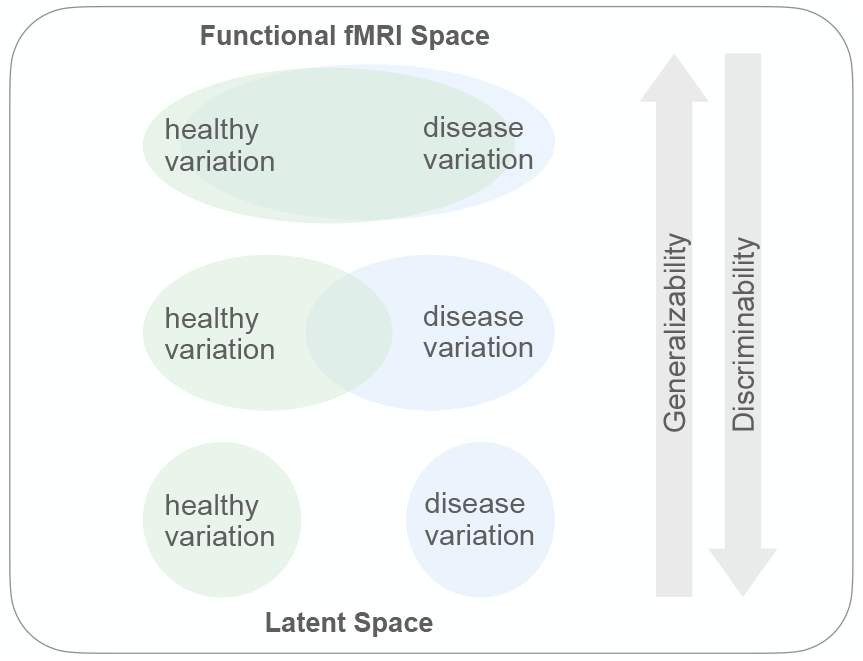
In the native functional fMRI space, there is a significant overlap between the variability of the BOLD signal observed in healthy and diseased patients. Dimensionality reduction methods such as PCA in this native space tend to capture this shared variance. Hence, the learned lower dimensional structure loses its discriminative power. Our metric learning framework allows one to map the functional data to a lower dimensional metric space (i.e., latent space), where one can tune the balance between discriminability (discriminative information in the embeddings), and generalizability (variance in the data captured by the embeddings).

To uncover latent low dimensional representations from fMRI volumes that improve inter-class variability while reducing intra-class variability, in this paper, we draw inspiration from related work in computer vision for face detection (Schroff et al., 2015) and more generally metric learning (Murphy, 2023). Face detection poses an analogous challenge to the one discussed above; face detection aims to learn face image representations for an individual such that they are invariant to environmental factors such as lighting and camera angle. In addition, these representations must also capture low-dimensional image features that allow for discrimination between a set of face images from two different individuals. The former goal in face detection is equivalent to minimizing intra-group variability within the same “context”, for example, within fMRI data from the same disease group. Additionally, the latter objective improves inter-group variability between “contexts”, for example, between fMRI data across different disease classes.

In this work, we extend our prior work (Mittal et al., 2022) and propose a metric learning framework that 1) generalizes the notion of contexts for different fMRI imaging paradigms, 2) leverages Information Metric Learning (IML) to learn lower dimensional embeddings from high dimensional fMRI data, and 3) uses advances in few-shot learning techniques to propose Prototypical Metric Learning (PML), a metric learning method for fMRI data when the number of contexts exceeds the number of samples per context. We demonstrate the use and efficacy of the metric learning framework by comparing Information Metric Learning (IML) and Prototypical Metric Learning (PML) against alternative methods on two publicly available fMRI datasets. Using IML and PML results in superior quantitative and qualitative outcomes on these datasets.

## 2. Methods

We propose a metric learning model that learns an informative latent representation of the data in a lower dimensional space. Given a spatiotemporal neural dataset 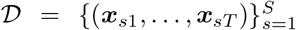, acquired from *S* subjects, where ***x***_*st*_ *∈* ℝ^*D*^ corresponds to an image at time *t* for subject *s*, we wish to encode an image ***x*** and its latent representation ***z*** in a way that maximally preserves the mutual information defined as:

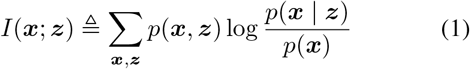

Where ***x*** *∈* ℝ^*D*^ and *∈* ***z*** ℝ^*K*^ correspond to an image and its lower dimensional latent representation at time *t* for subject *s*, minimizing the above information loss *I*(***x***; ***z***) by learning optimal latent representation ***z*** for each image ***x***, encourages the latent variables ***z*** to maximize the context information captured by the latent variables as described in the section below.

### 2.1 Definition of Context

It is important to establish that the “context” defined in this study can extend beyond its traditional boundaries. Beyond the conventional understanding of context in analyzing resting-state fMRI data, such as group condition or disease state, it encompasses scenarios like identifying the target event/block within a movie. This expanded perspective accommodates label-based analyses and opens avenues for event/block-based investigations, akin to the approach employed in resting state analyses. By acknowledging this flexible definition of context, we enable a comprehensive framework that caters to both label-driven and event-based analytical paradigms.

### 2.2 Information Metric Learning

Given an image ***x***, we wish to encode its latent representation ***z*** in a way that maximally preserves the mutual information *I*(***x***; ***z***). As it is computationally intractable to directly optimize the mutual information term *I*(***x***; ***z***), we optimize the InfoNCE loss (Oord et al., 2018) defined as:

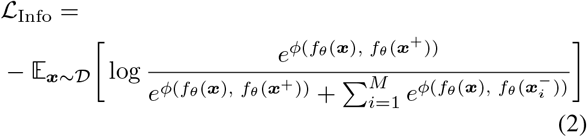

Where ***x, x***^+^*∈* 𝒟 ^+^ and 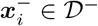 We define 𝒟^+^ to be the set of *positive* images, that is “in-context” with respect to a given image ***x***_*st*_, and 𝒟^*−*^ to be the set of *negative* images, that is, “out-of-context” with respect to ***x***_*st*_. For a labeled dataset, if the class of ***x***_*st*_ is *c*, then the image sets 𝒟^+^, 𝒟^*−*^ can simply be defined as:

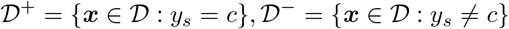

Here, *ϕ*(·, ·) can be any positive real-valued function, and *f*_*θ*_ : ℝ^*D*^ *→*ℝ^*K*^ is a function that maps a data point ***x*** from *D* dimensional observation image space to a latent vector ***z*** in the *K* dimensional latent space. Since, the InfoNCE loss lower bounds *I*(***x***; ***z***) (Oord et al., 2018; Poole et al., 2019):

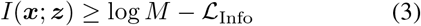

optimizing _Info_ objective maximizes the mutual information between the data points ***x*** and their respective latent representations ***z*** within the same context. Generally, one can choose the context to be defined by other variables such as age, gender, diagnosis, etc. However, it should be noted that contexts can also be defined in the absence of labels as we demonstrate in subsequent sections. For instance, in the case of event-related design for fMRI acquisition, one can use context variables for appropriate time locking for anchor, negative, and positive images.

### 2.3 Prototypical Metric Learning

Frequently observed within neuroimaging datasets is a scenario where the number of contexts significantly exceeds the available sample images for each context. For instance, there could be far more stimuli than the number of samples per stimulus. In such a setting, we modify the preceding approach and instead learn prototypes ***c***_1_ … ***c***_*K*_ for each context. Taking inspiration from prototypical networks (Snell et al., 2017), we propose prototypical information loss, which modifies the information loss discussed above and aims to learn latent prototypes ***c***_1_, …, ***c***_*K*_ for each of *K* contexts:

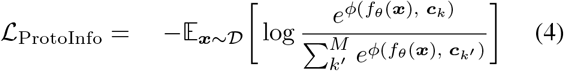

where a context vector ***c***_*k*_ is defined as:

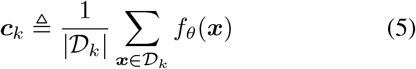

𝒟_*k*_ is the set of all data points that belong to the *k*-th context and *f*_*θ*_(***x***) is the mapping function (encoding neural network) of image ***x***. Note that 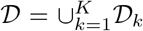 that is, all data points belong to at least one context.

### 2.4 Similarity Function and Latent Map

The information loss and the prototypical information loss are two metric learning approaches. However, for a specific problem instance, we must also define the similarity function *ϕ*(*·, ·*), and the mapping function *f*_*θ*_(·). Two common similarity measures are the cosine similarity:

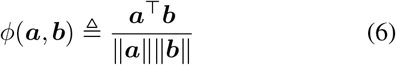

or the squared Euclidean distance:

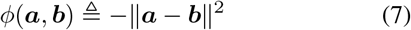

The choice of the similarity function affects the geometry of the underlying latent representation of that data. We choose the cosines distance when we only wish to model the angular distance between the representations and be invariant to the magnitude of the latent representation.

Additionally, we can choose *f*_*θ*_(·) to be a simple linear map:

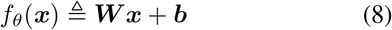

or a non-linear map parametrized by a neural network of the form:

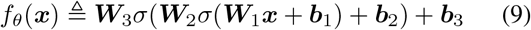

where *σ*(·) is a chosen non-linear function such as the softplus function. A non-linear map can help uncover a rich lower dimensional manifold underlying the data, whereas, a linear map can be particularly beneficial as it allows for a more interpretable mapping function and prevents overfitting particularly when ***x*** is very high dimensional.

To demonstrate specific instances for both approaches above, we use the following information loss with cosine similarity and a non-linear map for a sequence classification task on the ABIDE-150 dataset (Nielsen et al., 2013):

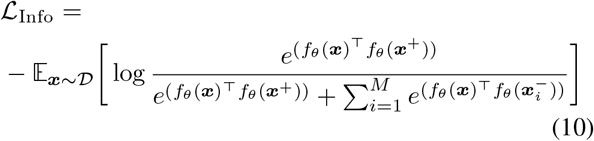

Where the outputs of the neural net *f*_*θ*_(·) are normalized to be unit norm.

For the segment recall task on the Raiders dataset (Haxby et al., 2011), we use the prototypical information loss with Euclidean distance and a linear map, resulting in the following loss objective:

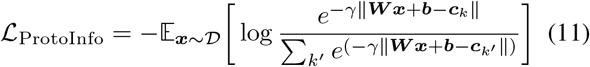

We describe these datasets in the Experiments section.

### 2.5 Model Inference

Model inference aims to optimize the information losses discussed above or equivalently maximize the lower bound to the mutual information *I*(***x***; ***z***). The optimization objective then is to learn the optimal latent representation ***z***^*∗*^ for each image ***x*** *∈* 𝒟:

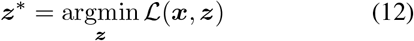

This objective is parameterized with a map *f*_*θ*_(·), with learnable parameters *θ*, where ***z*** ≜ *f*_*θ*_(***x***). Hence, inference is amortized through the neural network *f*_*θ*_(·), this allows for fast inference without the need for solving a new optimization problem for each unseen data point ***x***. We optimize the parameterized objective with respect to the parameters *θ*:

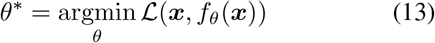

where *θ* corresponds to the parameters of the mapping function: weights and biases of the neural network or the matrix ***W*** in the case of a linear map.

We can compute the gradient of this parameterized objective with respect to the parameters *θ* :

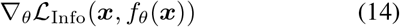

This loss objective can be easily optimized using standard neural network stochastic optimization methods such as gradient descent with ADAM (Kingma & Ba, 2014).

The training algorithms for specific instances of both information metric learning and prototypical metric learning are outlined in **Algorithm 1** and **Algorithm 2**.

#### Algorithm 1

Information Metric Learning

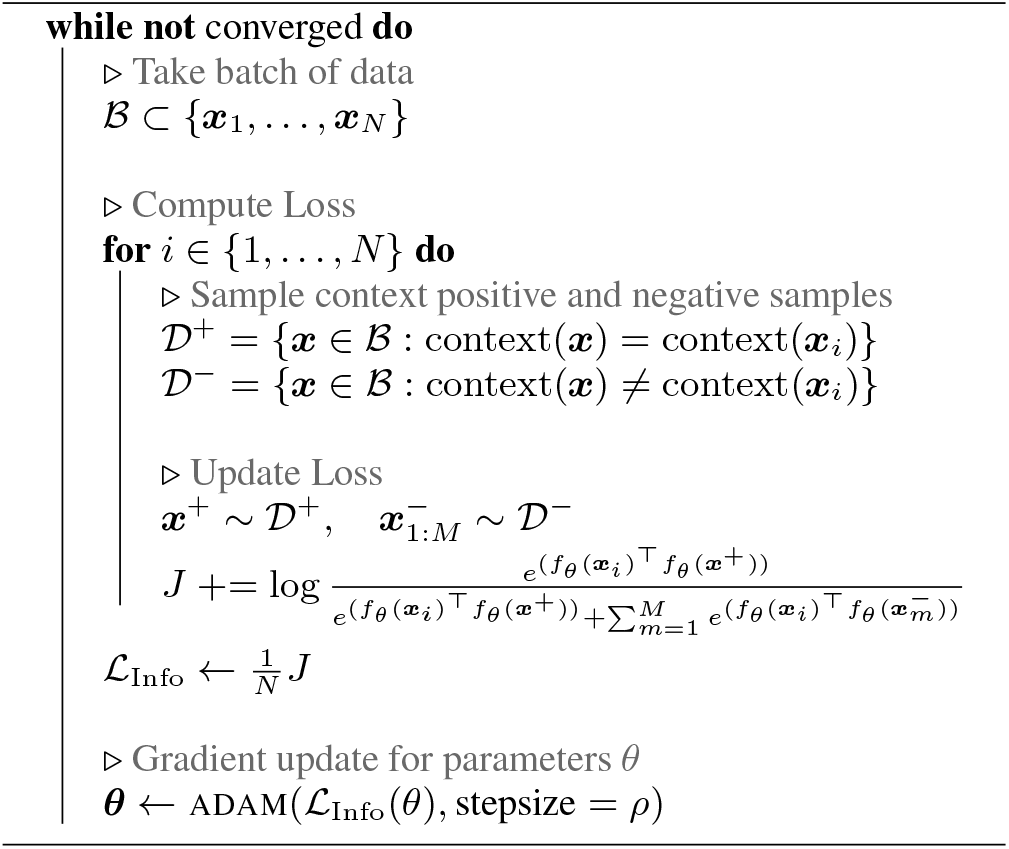

#### Algorithm 2

Prototypical Metric Learning

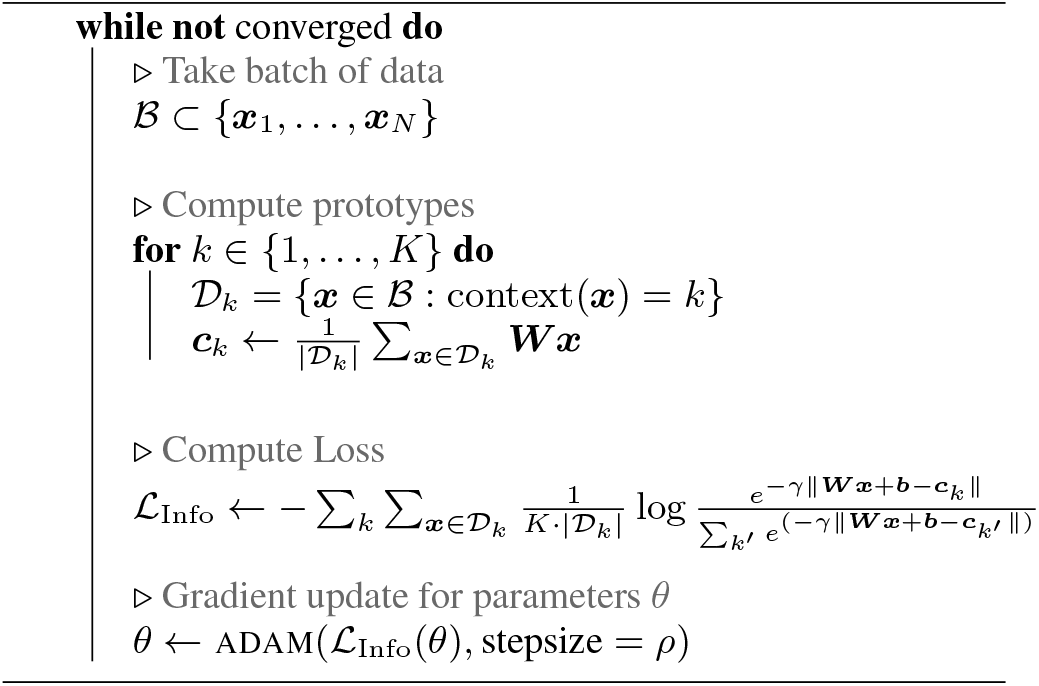

The context(·) : ℝ^*D*^ *→*ℤ^+^ function is a bespoke function that maps any data point to a context. Trivially, it could be the function that maps data points to its provided labels for a labeled dataset. However, alternative context functions allow for richer representation learning as we discuss in the experiments section.

### 2.6 Related Work Used for Comparison

In this section, two related methodologies – Graph Convolutional Network (GCN) and Principal Component Analysis (PCA) – are introduced. Below (in the Experiments and Results section) we conduct a comparative analysis obtained from these two methods as a way to establish the effectiveness of our proposed IML and PML approaches.

#### 2.6.1 Graph Convolutional Network

Ktena et al. proposed a graph convolutional neural network (GCN) for learning representations of brain connectivity networks (Ktena et al., 2018).

GCN utilizes a graph neural network *h*_*ϕ*_(·), which takes covariance matrix computed from a subject’s time series sequence (***x***_1_, …, ***x***_*T*_) as the input, and outputs a latent embedding ***z***_*i*_. Subsequently, GCN uses a modified Global Loss objective proposed by Kumar BG et al., which seeks to minimize the distance between “in-context” representations of images, while also maximizing the distance between “out-of-context” latent representations of images (Kumar BG et al., 2016). The loss function proposed for GCN can be written as (Ktena et al., 2018):

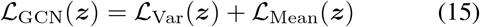

Where, the variance loss ℒ_Var_ leads to the minimization of the variance between all the “in-context” and “out-of-context” latent vectors:

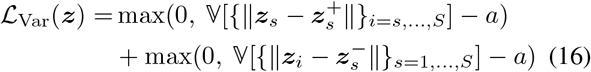

whereas the mean loss ℒ_Mean_, minimizes the expected distance between “in-context” latent vectors, while maximizing the distance between “out-of-context” latent vectors.

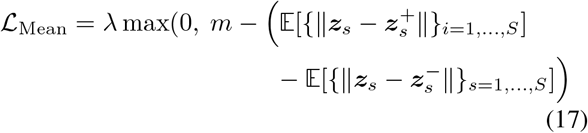

where ***z***_*i*_ is the embedding for the *i*-th subject sequence, and ***z***^+^ *∈* 𝒟 ^+^, ***z***^*−*^ *∈* 𝒟^*−*^ are the positive and negative samples from the positive and negative context image sets, described in the above sections. The latent variables are learned through the neural network:

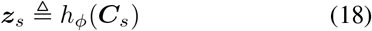

where ***C***_*s*_ is the subject covariance matrix computed using the subject *s*’s image sequence (***x***_*s*1_, …, ***x***_*sT*_).

#### 2.6.2 Principal Component Analysis

Given the widespread use of PCA in neuroimaging literature as a tool for dimensionality reduction, we compare our method to PCA as well. We can write the generative model for PCA (Bishop, 2006) as:

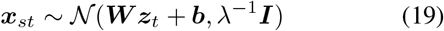

which corresponds to the loss function:

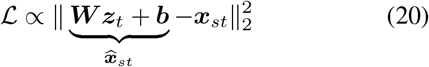

where the loss objective encourages the latent variable to capture variance in the observed data by minimizing the reconstruction loss between the observed data ***x***_*st*_ and the reconstructed data 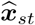 Here the class or context information is not explicitly utilized to learn the latent structure and hence subtle variations in data that are informative for discriminating contexts might be lost to underlying dimensions of variation that vary significantly despite not being informative for a given context.

## 3. Experiments

We evaluate our proposed method on ABIDE-150 (Nielsen et al., 2013), a resting state fMRI dataset of healthy subjects and people with autism, and the Raider’s dataset (Haxby et al., 2011), a dataset composed of fMRI volumes acquired while multiple subjects watched the same movie. In the following sections, we describe the datasets, and our evaluation task, and compare our results against a simple PCA+kNN approach, as well as the current state-of-the-art results on the ABIDE-150 dataset achieved by GCN.

### 3.1 Datasets

#### ABIDE-150

Autism Brain Imaging Data Exchange is a multi-site resting-state fMRI dataset comprising scans from healthy subjects and clinically diagnosed subjects with autism (Nielsen et al., 2013). Our experiments use a 150-subject random subset (autism=74, control=76) from this dataset, restricted to the NYU imaging site. Each subject has a sequence length of 78 volumes (TRs). The images were pre-processed using C-PAC pipeline (Craddock et al., 2013) without global signal regression.

#### Raiders Dataset

The Raiders dataset (Haxby et al., 2011) consists of fMRI scans from 10 subjects that were collected while subjects viewed the movie “Raiders of the Lost Ark” (110 mins). We used a pre-processed version of the dataset (Kumar et al., 2020) containing 1000 voxels (500 per hemisphere) from the v,entral temporal (VT) cortex. Each TR was 3s long, with *≈*2200 TRs in total for a 110-minute movie.

### 3.2 Evaluation Tasks

#### Disease Sequence Classification

We compare Information Metric Learning against GCN (Ktena et al., 2018) on the ABIDE dataset for a subject classification task. The goal is to learn latent representations of data, which can then be used to classify the sequences into disease and healthy classes. In our work, instead of using the covariance matrix, we infer each volume’s lower dimensional latent structure and disregard the temporal structure between consecutive volumes. To classify a sequence, we use k-nearest neighbors to label each volume in a sequence and then use a majority label across all volumes in a sequence to infer the class label of the entire sequence.

#### Segment Matching

We evaluate the latent representation of the acquired fMRI sequence by testing if we can use the inferred latent representation to identify the correct 30-second (10 TR) time segment of a movie that a held-out subject was watching. That is, given sequences of fMRI activity from 9 subjects watching the same movie, can we identify where in the sequence a 30-second time segment for the held-out 10th subject belongs? Alternatively, given a sequence of fMRI activity segments corresponding to VT activity in 9 subjects watching a movie, if we are given a randomly permuted sequence of fMRI activity segments, how many segments of the 10th subject can we correctly re-order to the correct sequential order?

### 3.3 Experimental Details

#### Information Metric Learning on ABIDE-150

We evaluate information metric learning against GCN and PCA on the ABIDE dataset. As a first step for dimensionality reduction, instead of using predefined anatomical atlas, we use dictionary learning (Mairal et al., 2009) to learn a functional atlas composed of regions of interest (columns of the dictionary matrix). The dictionary components define the 80 ROIs for our data. As a first step, we map all sequences from our dataset to this 80-dimensional space, which is then used to train and evaluate all models.

To compute the PCA benchmark, we calculate the first ten principal components (PCs) from the training data and then project the held-out data onto these 10 PCs, sorted by captured variance. We project each high-dimensional fMRI time series to an 80-dimensional time series, which is then projected to a 10-dimensional time series. We then use a K-nearest neighbor classifier (K-NN) with *K* = 3 to label each volume of each projected held-out sequence; we then take the majority of all the labels in a sequence to assign the final class label to a held-out sequence. We repeat this for all held-out rs-fMRI sequences to classify all the held-out sequences. We repeat this process across five cross-validation folds and report the results in Table 1.

**Table 1.**
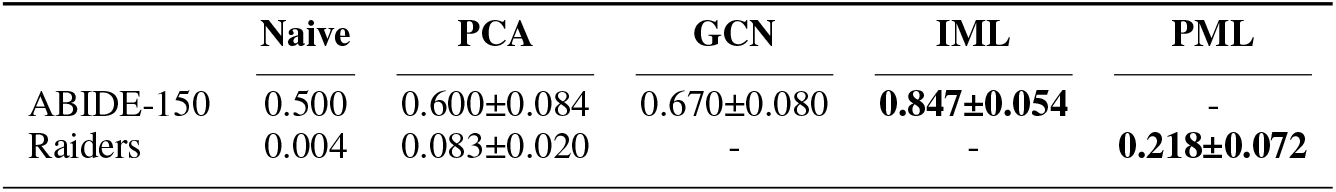
We report the mean and standard deviation of 5-fold cross-validation accuracy across the ABIDE-150 and Raiders datasets. For both datasets, the naive method randomly assigns context to held-out data to give us a trivial baseline for comparison. IML significantly outperforms the naive and PCA methods, as well as the current state-of-the-art GCN method. PML significantly outperforms the Naive and PCA baseline performance.

|  | Naive | PCA | GCN | IML | PML |
| --- | --- | --- | --- | --- | --- |
| ABIDE-150 | 0.500 | 0.600±0.084 | 0.670±0.080 | **0.847±0.054** | - |
| Raiders | 0.004 | 0.083±0.020 | - | - | **0.218±0.072** |

For information metric learning on the ABIDE dataset, we treat each of the volumes across all subjects 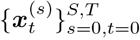 as distinct anchor points (see Figure 2). Since labels are provided for the training this data set, we define the context(·) function as:

**Figure 2.**
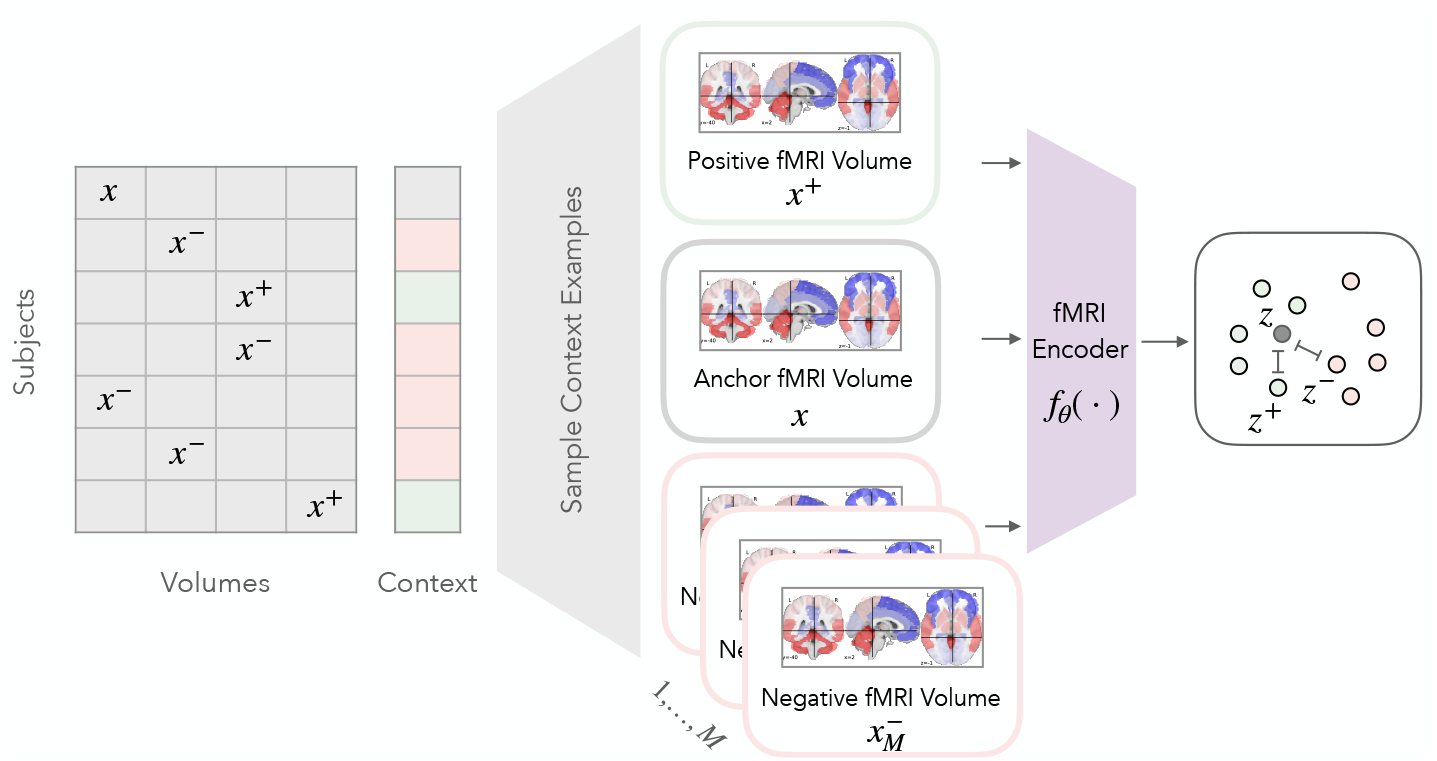
In information metric learning (IML), the anchor fMRI volume, the positive fMRI volume, and the negative fMRI volumes are sampled across subjects and the volume sequence. Each volume is then mapped through an encoding neural network *f*_*θ*_(·) to a lower dimensional space. The information metric learning loss encourages fMRI volumes belonging to the same context, in this case, disease or healthy, to be embedded close together in the latent space. In contrast, those from different contexts are embedded further apart. Since the newly learned space is a metric space, we can use popular modeling methods such as density modeling with mixture models in this new space.

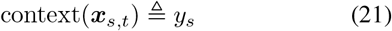

As discussed in the methods section, we use cosine similarity, with a 3 layered neural network as the map function of the form:

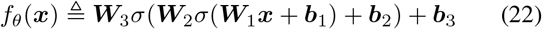

with ***W***_1_ *∈* ℝ^80*×*50^, ***W***_2_ *∈* ℝ^50*×*20^, and ***W***_2_ *∈* ℝ^20*×*10^. With *σ*(·) ≜ Softplus(·) non-linearity. Additionally, to prevent overfitting, we add dropout layers between each of the layers with a dropout probability of 0.3. For the loss term, we choose *M* = 20. Finally, to train this network, we use ADAM optimizer with learning rate = 0.001, betas=(0.9, 0.999), eps= 1e*−* 08, weight decay= 0.0. We train the model for 40 epochs, where each batch contains the data across 30 random subjects. To evaluate the accuracy of the held-out subject set, we use a K-nearest neighbor classifier to classify each volume and then take the majority class across all volumes in a sequence to assign the final class label to the sequence. Then, the metric learning algorithm (Algorithm 1) learns to embed volumes that belong to the same disease class, close together in the latent space, and volumes from different classes far apart. Metric learning with this context function learns embeddings for subjects while maximizing invariance across a disease class.

#### Prototypical Information Metric Learning on Raiders

To compute the PCA benchmark on the Raiders dataset, we first reduce the dimensionality of the fMRI volume sequences by projecting each volume (1000 voxels) onto the first ten principal components, sorted by variance captured. Then, for each contiguous sequence of 10 TRs (volumes), we concatenate the projected 10-dimensional vectors to form a 100-dimensional vector. Hence, the sequence of 2203 volumes (TR=3s), corresponding to the 110-minute movie runtime for each subject, is reduced to 219 blocks of 100-dimensional segments for each subject. We then use a K-nearest neighbor classifier with *K* = 1 to assign the temporal sequence to each held-out block. We compute accuracy as the percentage of held-out blocks with correctly assigned temporal sequences.

For prototypical metric learning (see Figure 3), as a first step of dimensionality reduction, we perform the same steps for dimensionality reduction as described in the PCA dimensionality reduction, except we project the data onto 20 principal components. Compared to the PCA benchmark, we start with twice the dimensionality, allowing our algorithm to uncover a lower-dimensional structure. To apply the prototypical metric learning framework, we define the context(·) function as:

**Figure 3.**
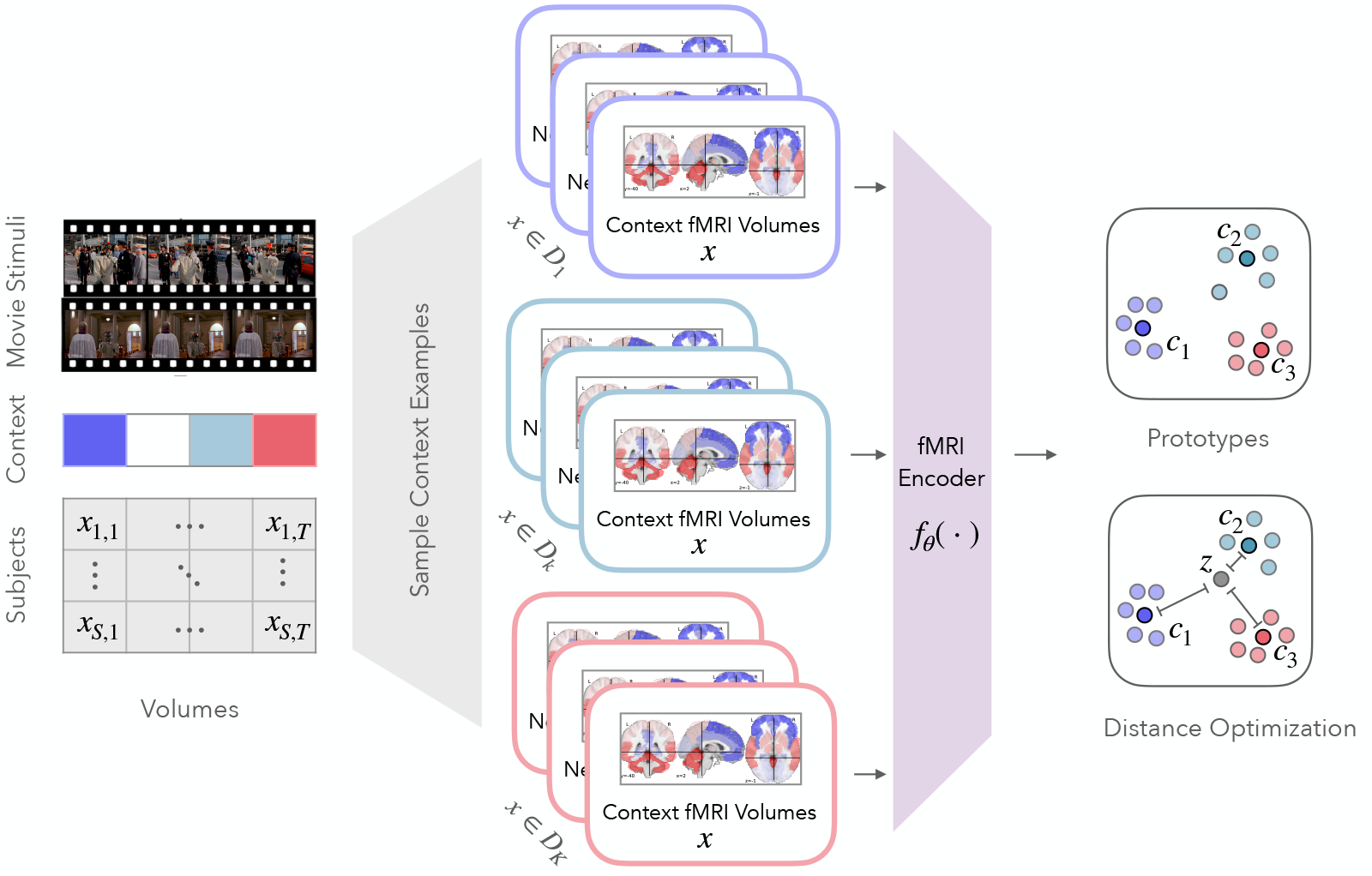
In prototypical metric learning (PML), all the fMRI volumes belonging to a particular context *D*_*k*_, in this case, the contexts, corresponding to movie blocks, are encoded through the fMRI encoding model. The encoded embeddings are then used to compute prototypes, which are the latent centroids for each context. The optimization process then optimizes the distances between the embeddings such that each embedding moves towards its own context prototype while away from other context prototypes.

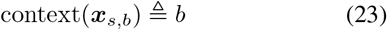

Hence, there are 219 contexts *{****c***_1_, …, ***c***_219_ *}*. Note that held-out contexts are computed from held-out data. For our mapping function, to prevent overfitting we use a simple linear mapping function:

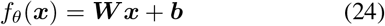

with ***W*** *∈* ℝ^200*×*20^. The function maps the PCA reduced and concatenated 200-dimensional vector to a 20-dimensional embedding vector. As discussed in the methods section, we use the squared Euclidean distance for this dataset. We train the model using the ADAM optimizer with the same learning rate parameters as in the ABIDE dataset. However, we use full batch optimization steps to ensure high-quality context vectors are generated in each epoch of the optimization process. We train for a total of 800 epochs. Finally, to assign the correct sequence position to a given block ***x*** using one nearest context classifier:

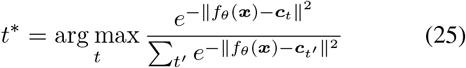

The prototypical metric learning algorithm (Algorithm 2), for the Raiders dataset learns a lower dimensional representation that captures only the variance that is specific to the temporal visual stimulation effect of movie scenes while maximizing invariance to all other inter and intra-subject variation.

For both evaluation cases above, there is no overlap of subjects across the training set and the held-out sets. This makes the task significantly more difficult as the models need to account for variations in subject-specific anatomy, physiological differences, scanner differences, etc. Note that the context function does not require labels but is defined to maximize invariance across a dataset bespoke context. For instance, one could define a forward-looking context function for contrastive learning context(***x***_*s,t*_) ≜ 𝕀[*t < k*]. We do not discuss contrastive learning applications in this work and leave it as a future area of exploration.

## 4. Results

We report the 5-fold cross-validation accuracy in Table 1 where the accuracy for the ABIDE-150 dataset corresponds to the percentage sequences correctly classified as disease vs. healthy, whereas the accuracy for Raider’s dataset corresponds to the models’ accuracy of correctly identifying sequential positions of all the blocks from an unseen heldout subject sequence.

In our experiments, information metric learning outperformed PCA as well as the Graph Convolutions Network proposed by Ktena et al. on the ABIDE-150 dataset sequence classification task by a considerable margin. Demonstrating that information metric learning can learn a map function to an underlying low dimensional space in data that generalizes across subjects better than latent representations learned using PCA or GCN. Prototypical Metric Learning had a 2.5x performance improvement over PCA despite PML only using 20-dimensional embeddings for each time block as compared to 200-dimensional embeddings used by PCA.

We examine the learned latent embeddings for IML and PCA using Umap (Sainburg et al., 2021) (Figure 4A), where each point corresponds to the latent embedding of an fMRI volume. IML learns an underlying non-linear manifold that captures the variance of the data while improving the discriminability between contexts. For the interpretation of the original fMRI volumes, we found the anterior-posterior underconnectivity in the Autism patients as reported in other literature (Cherkassky et al., 2006; Heinsfeld et al., 2018). Latent embedding also captures the BOLD difference in the occipital cortex/vision network between healthy and autistic brain (McKinnon et al., 2019)(see **Top left** and **Bottom left** of Figure 4A). PCA, on the other hand, optimizes the variance captured by the latent embeddings. However, since the embeddings learned using PCA maximize the variance captured and the principal components might be misaligned with the axis that allows for context discrimination, useful information is lost during dimensionality reduction.

**Figure 4.**
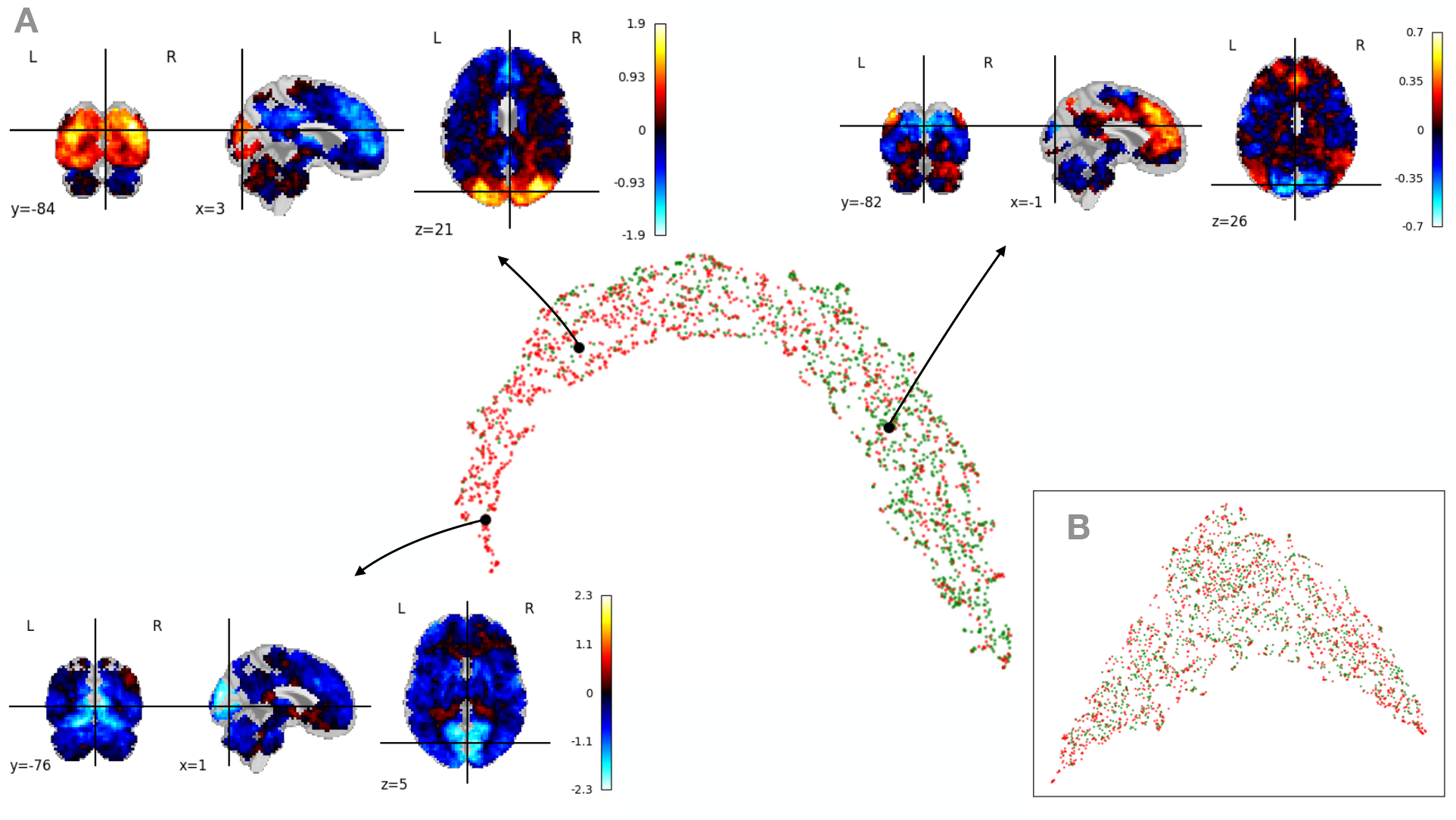
**A**: Umap visualization for the learned lower dimensional latent space on the ABIDE-150 dataset. Each point corresponds to the latent embedding of fMRI volumes from healthy subjects (green) and people with autism (red). The information metric learning model learns a lower dimensional manifold, which captures the data’s variance while mapping embeddings from the same context closer together. Additionally, the original fMRI volumes of the mapped embeddings are interpretable. Several images of fMRI volumes were illustrated as an example. **Top left**: fMRI volume figure shows the anterior-posterior anti-correlation observed in the autistic brain (Cherkassky et al., 2006; Heinsfeld et al., 2018). **Bottom left**: an example supports the evidence from the literature that there is a reduced resting-state BOLD within the occipital cortex/vision network in Autism (McKinnon et al., 2019). **Top right**: a counterexample of top left observed in healthy subjects. **B**: Umap visualization for latent space learned using PCA, which uncovers a lower dimensional manifold that captures the variance in data. However, since PCA does not utilize context information, it reduces discriminability between the latent embeddings.

tTo compare the aggregate performance of PML against PCA, we compute the kernel density estimate over the probabilities that the latent embeddings corresponding to any time block are closest to the centroid for that time block (Figure 5). We note that relative to embeddings learned through PCA, the vast majority of embeddings learned using PML are mapped closer to their corresponding centroids, illustrating that PML, even with a linear map, can learn structure that is informative of context.

**Figure 5.**
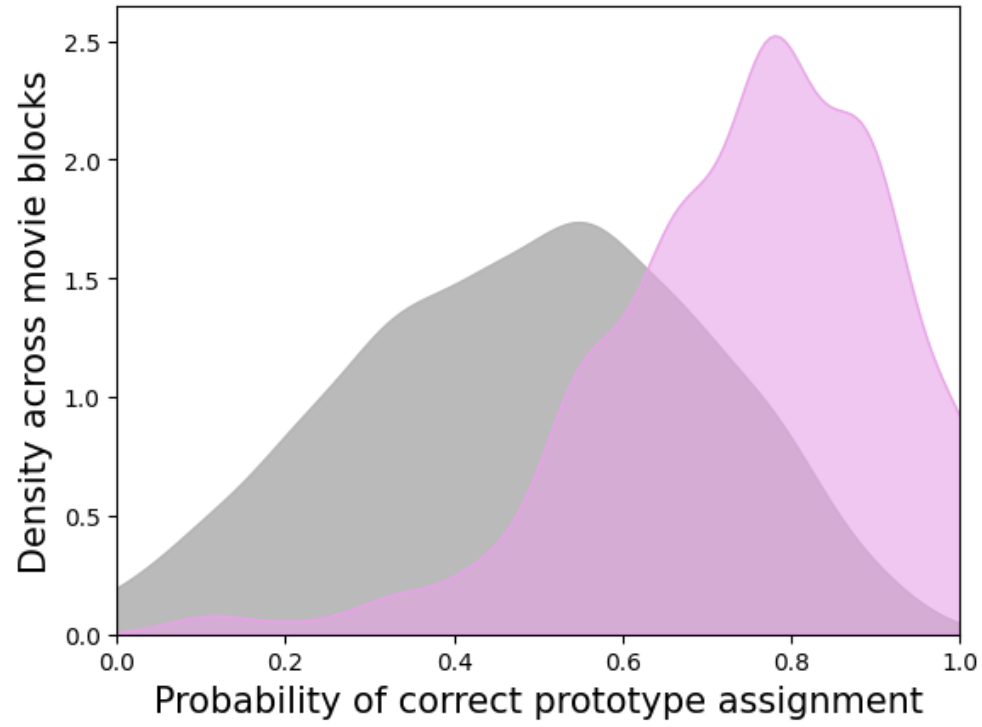
Kernel density across all time blocks of the Raiders dataset, where the x-axis corresponds to the probability of an fMRI volume embedding being correctly assigned to its closest context centroid. Embeddings learned using information metric learning (pink) have a far higher probability of being correctly assigned compared to the embeddings learned using PCA (gray).

In PML, the *γ* parameter controls the uncertainty of the model (Figure 7). Larger values of *γ* encourage higher uncertainty, while smaller values of *γ* encourage sharp probability. This is useful for certain datasets like the Raiders dataset as two movie blocks can be very perceptually similar. We would expect very similar responses in the visual cortex, so we expect the embeddings corresponding to both time blocks to be close to the same prototype. Adjust *γ* allows for “slack” in the model and allows us to model this uncertainty.

To approximate the variance of the data captured for each context, we calculate the z-scored distance of each latent vector belonging to a context from the mean latent representation corresponding to that context:

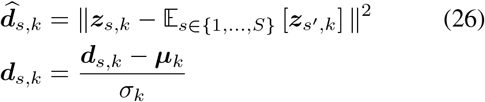

We compare these distances across the raw bold data, embeddings learned using PCA, and embeddings learned using PML (Figure 6). In the case of raw bold data ***z***_*s,k*_ is simply ***x***_*s,k*_. As desired, the learned variance captured by both PCA and PML closely follows the variance in the raw bold data. However, PML representations reweight the variance captured also to be informative of the context.

**Figure 6.**
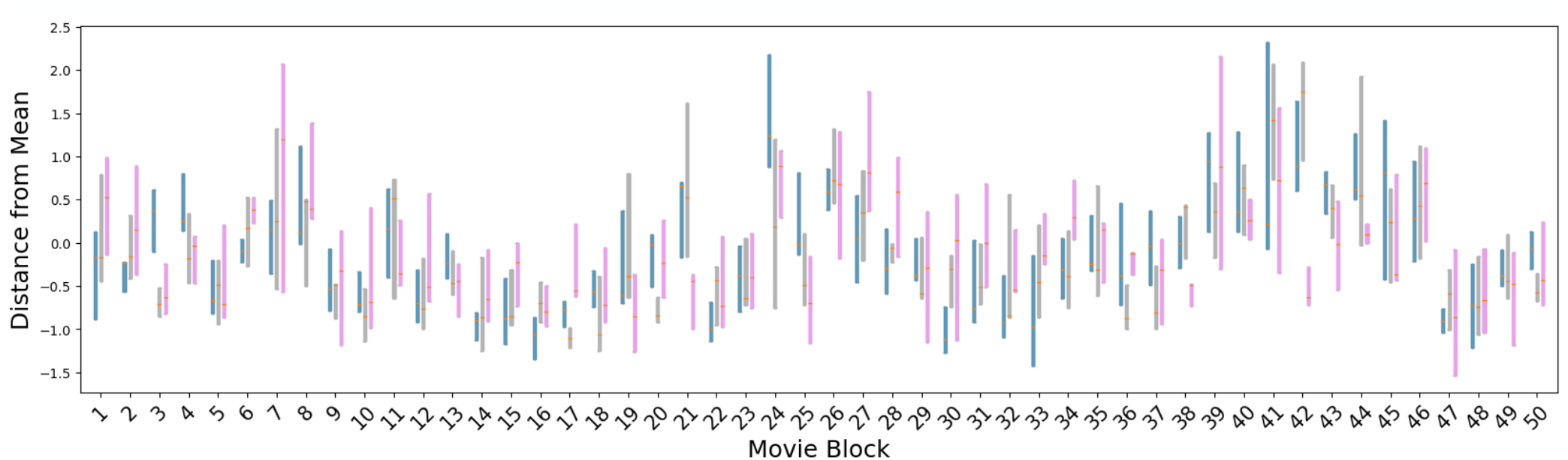
The 95% confidence interval for z-scored distances of data belonging to a time block from the centroid computed for that time block for the first 50 time blocks of the Raiders movie. The results from raw bold data (green), PCA (gray), and PML (pink) are shown. We note that both PCA and PML capture the variance across each time block. However, PML trades off some variance (closer to 0) to learn representations that are also informative of the context.

**Figure 7.**
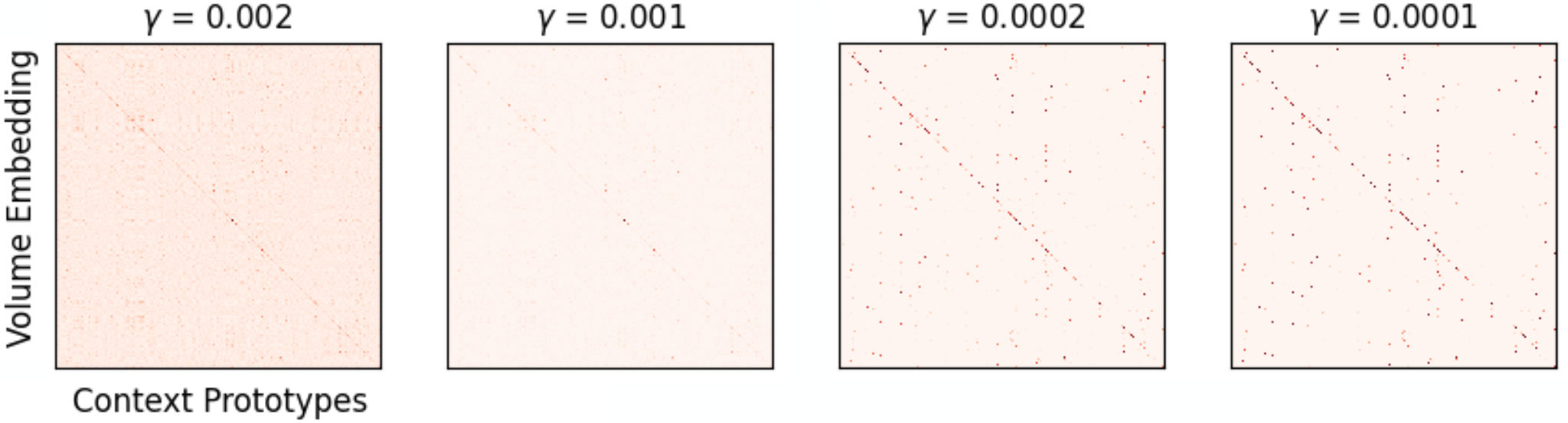
Each matrix here shows the probability of assignment of each volume (rows) embedding to one of the 219 prototypes (columns). All matrices share the same x and y label. In prototypical metric learning, the *γ* parameter controls the uncertainty in our predictions. For smaller values of *γ*, the model has sharper predictions, while for larger values of *γ*, the probabilities are diffused over multiple possible outcomes (*γ* decreases from left to right).

## 5. Discussion

One can interpret the IML and PML methods as two-step processes combined into one–i.e., a dimensionality reduction followed by a clustering. Our metric learning approach learns to map high dimensional data into low dimensional space such that the points from the same context cluster together. In the case of IML this objective is implicit. That is, since points belonging to the same context are pulled closer together while points from different contexts are pushed farther apart, this, in effect, clusters the points from the same contexts together. PML’s clustering process is more explicit as prototypes for each context are explicitly computed. Then, the distance between each embedding point belonging to a context and the prototype for that context is explicitly minimized. At the same time, the distance of these points from the prototypes of different contexts is maximized.

PCA can be viewed as a generative model where the objective is to minimize the reconstruction error. The latent are then learned so that the data can be best reconstructed (Bishop, 2006). This task is challenging for high dimensional data with a modest number of samples, as this might involve the latent modeling information in the data at fidelity that is not required by the downstream task. For example, individual voxel differences are often not informative or robust enough for generalization across a population. Metric learning provides a middle ground between generative and discriminative learning. The goal changes from learning a latent representation that can be used to reconstruct the data to the goal of learning a generalizable latent representation that allows discriminability of contexts. Learning a distribution *p*(***z*** |***x***) is a more modest goal as it does not require us to model a high dimensional image distribution *p*(***x***|***z***).

Popular non-linear extensions of PCA such as the Variational AutoEncoder (VAE) (Kingma & Welling, 2013; Dilokthanakul et al., 2016) model both *p*(***x*** |***z***), the decoder, and *p*(***z*** | ***x***), the encoder. In IML and PML, we only model *p*(***z*** | ***x***). The signal that informs the encoding distribution is not the reconstruction error but the metric loss *p*(context |***z***). IML and PML can further be extended to relax assumptions around the latent distribution. For instance, the magnet loss (Rippel et al., 2015), allows us to model multimodal clusters.

Other supervised autoencoders such as Random Forest (RF) (Ho, 1995) and Support Vector Machine (SVM) (Vapnik, 1998) have also been introduced where they aim to maximize the width of the gap between categories. More recently, deep neural networks (DNN) have also been used in the classification analysis of rs-fMRI data (Abraham et al., 2014; Khosla et al., 2019). However, the results of those methods on ABIDE dataset are not as competitive as ours from IML (IML: 84.7%, DNN: 70%, RF: 63%, SVM:65% (Heinsfeld et al., 2018)).

In conclusion, our metric learning framework (IML and PML) enables a reduction of high-dimensional neuroimaging data into meaningful latent low-dimensional structures linked to behavior, disease, or in an experimental context. Unlike more traditional approaches, such as PCA, these methods learn the latent embeddings that reduce the intra-group variability while maximizing the inter-group variability. We show results on fMRI datasets that demonstrate superior performance to well-used alternative approaches. We also provide Python code and examples so that other researchers can use these methods on their data.

## 6. Codes and Materials Availability

All presented algorithms are implemented in Python 3.7. Codes and example results are available at Github.

## 7. Acknowledgements

This work was supported by a Vannevar Bush Faculty Fellowship from the US Department of Defense (N00014-20-1-2027) and a Center of Excellence grant from the Air Force Office of Scientific Research (FA9550-22-1-0337).

